# Innovative Prevention Approach for Pan Allergens-Induced Allergy: Effective Regulation of Heat Shock Protein 70 and Profilin 3 by siRNA-Loaded Nanomaterial

**DOI:** 10.64898/2025.12.09.693215

**Authors:** Zhiqi Liang, Bohan Ren, Kexin Yan, Peijiao Zhang, Sophia Zheng

## Abstract

Currently, the widely accepted approach to addressing pollen allergy primarily through altering the immune system of patients. However, the great potential of genetic engineering for downregulating allergenic proteins and reducing allergenicity is often overlooked in previous studies. Here, we report an effective solution to reducing plant allergenicity by utilizing RNA interference (RNAi) technology to calibrate levels of pan-allergenic proteins Heat Shock Protein 70 (HSP 70) and Profilin 3 (PFN 3) in transgenic Arabidopsis, Tobacco, and Osmanthus. This has been shown in research to effectively mitigate and prevent allergenic reactions in humans. 21-bp long siRNA sequences encased by Polyethyleneimine modified carbon dots (PEI-CDs) are synthesized to constitute a nanoparticle-mediated delivery system to efficiently deliver small-interfering RNA (siRNA) vectors into plants. Afterwards, RNA was extracted from *Osmanthus fragrans* and *Gardenia jasminoides* and underwent reverse transcription quantitative-polymerase chain reaction (RT-qPCR). The result indicated the expected suppression of mRNA corresponding to the targeted proteins, proving the functional aspect of RNAi in allergen suppression trunk injection and suggesting its applicability in real-life scenarios.

## 1 Introduction

Allergenic diseases represent a growing global health burden, affecting the quality of life for millions of individuals worldwide. Among these, IgE-mediated allergic reactions are one of the most common types of reaction. The binding of otherwise harmless exogenous proteins, i.e., allergens, with hypersensitive IgE antibodies triggers the release of histamine in mast cells, causing a reaction called allergy (Sullivan & Kushnir, 2005). Such responses induce symptoms ranging from mild dermatitis to lethal asphyxia. Abundant measures have been established to eliminate or mitigate this inappropriate immune response, the safest procedure being avoidance from potential allergy triggers.

Specifically, pollen allergy issues is a significant public health issue. According to the World Allergy Organization, between 10% to 30% of adults and up to 40% of children worldwide are affected by various allergic conditions (Sullivan & Kushnir, 2005). The increasing incidence of pollen allergies also poses substantial economic implications, as in the United States alone, over $3 billion is spent on pollen allergy medications annually, with more than half of this expenditure attributed to prescription drugs (CDC, 2024). The situation is further complicated by the effects of climate change. Between 1995 and 2011, rising temperatures in the United States resulted in an extension of pollen seasons by 11 to 27 days (Asthma & Allergy Foundation of America, n.d.). Warmer weather conditions prompt earlier blooming and prolonged pollen release from plants. Additionally, increased carbon dioxide levels stimulate greater pollen production, while regions that previously experienced colder climates are now becoming suitable for pollen-producing plants, further expanding the geographical range of allergens (Zhang & Steiner, 2022). Given this context, there is a pressing demand for innovative approaches to mitigate pollen allergies at the source.

As a result, we decided to focus on pollen allergy issues, particularly those caused by *Populus tomentosa*, especially in urban areas such as Beijing. *Populus tomentosa*, more commonly known as the Chinese white poplar, is extensively cultivated in northern China for its rapid growth and utility in urban landscaping. However, this widespread cultivation has led to increased pollen exposure during the spring season, when these trees release substantial amounts of pollen into the atmosphere. Statistically, of the approximately 21.84 million permanent population, it is estimated that at least 2 million residents—roughly 9.2%—suffer from pollen allergies, and thus it is recognized as a dominant spring aeroallergen. Epidemiological studies have consistently shown that its pollen is a primary trigger for seasonal allergic rhinitis and asthma exacerbations in the population during March and April (Wang et al, 2018). This pollen is recognized as a major aeroallergen, contributing to respiratory allergic diseases among a growing number of individuals sensitive to allergens.

To further corroborate this conclusion, a cross-sectional pathophysiological study reported differential allergenic pollen prevalence in major regions of China, and a remarkable abundance of Populus-related allergen cases was found in nearly all districts assessed, highlighting it as a strong candidate for our study.

**Table 1.** Predominant allergenic pollen types in different regions in China. (Yang et.al, 2015)

| Region | Main pollens allergens in spring | Main pollen allergens in summer and autumn |
| --- | --- | --- |
| Eastern China | *Pinus, Populus, Platanus, Broussonetia* | *Poaceae, Chenopodiaceae, Ambrosia artemisiifolia, Artemisia, Humulus japonicus* |
| South China | *Pinus, Cupressus, Moraceae, Broussonetia* | *Poaccae, Casuarina, Artemisia, Chenopodiaceae* |
| Central China | *Platanus, Pinus, Cupressus, Broussonetia, Populus* | *Artemisia, Poaccae, Ligustrum, Chenopodiaceae* |
| North China | *Populus, Cupressus, Betula, Pinus* | *Artemisia, Humulus japonicus, Chenopodiaceae, Poaccae* |
| Northwestern China | *Populus, Betula, Salix, Platanus* | *Artemisia, Chenopodiaceae, Humulus japonicus, Poaccae* |
| Southwestern China | *Salix, Populus, Cupressus, Platanus* | *Poaccae, Artemisia, Chenopodiaceae* |
| Northeast China | *Populus, Pinus, Ulmus, Betula* | *Artemisia, Poaccae, Humulus japonicus, Chenopodiaceae* |

Given the widespread prevalence of Populus pollen allergens across various regions in China, it is crucial to investigate the molecular nature of these allergens to identify key targets for allergy mitigation. Previous proteomic studies on poplar pollen have identified a range of potential allergens. A comprehensive review of this literature reveals that at least 28 proteins from Populus species have been characterized or predicted as allergens, belonging to 10 protein families. Among these, we selected two target proteins, Potri.010G205700.1 from the HSP 70 protein family and Potri.003G047700.1 from the Profilin family, for further evaluation based on multiple criteria.

First, we examined the prevalence of allergies activated by the protein. 2-Dimensional gel electrophoresis results from past literature have recognized the presence of Potri.003G047700 in mature *Arabidopsis* pollen, *Populus tomentosa* pollen, and tomato pollen tissues, demonstrating the targeted protein exists in many different plant species. A BLAST analysis conducted for the putative allergen Potri.003G047700.1 in the mature poplar pollen yielded a bit score of 411 (surpassed 62 other proteins in a group of 74 proteins observed), representing a high homology between this protein with other known allergens in the database, signifying a higher likelihood of eliciting allergenic hypersensitivity. Complementarily, the E-value of the protein was at 5.8 * 10^-37, a statistical validation of the significance of this inference (Zhang et al., 2015)

Similarly, BLAST analysis for Potri. 010G205700.1 was executed, receiving a bit-score of 605.2 and an E-value of 3.7 * 10^-174. The allergenicity of the proteins can be further confirmed a >35% sequence similarity to reported allergens, as well as the presence of ≥8 consecutive amino acids corresponding to previously identified allergens were found according to current information on the Structural Database of Allergenic Proteins and Allergen Online (Wang et al., 2018). Overall, the two proteins can be categorized as pan-allergens, a group of ubiquitous proteins, therefore widely found in many unrelated species (Hauser et.al, 2010). Thus, targeting these specific proteins is justified, as a strategy aimed at them could reduce the allergenicity of pollen from multiple plant species and extend the goals of Wang et al. to develop novel methods against these allergens.

To better understand their role in allergic reactions, it is important to explore the biological functions and allergenic potential of HSP 70 and Profilin 3.

HSP 70 is a member of the heat shock protein family, which is crucial for protein folding and cellular stress responses. It functions as a molecular chaperone, helping to maintain protein homeostasis under stress conditions. In allergic individuals, HSP 70 can induce IgE-mediated hypersensitivity reactions and activate T-cell responses. Studies have shown that HSP 70 is widely present in plant pollen, contributing to allergenic cross-reactivity between different pollen types and even between plant-derived foods. Elevated levels of HSP 70 have been detected in patients suffering from allergic rhinitis, indicating its role as a significant mediator in allergic reactions. Specifically, HSP 70 can bind to IgE antibodies in sensitized individuals, leading to the activation of immune pathways that exacerbate allergic symptoms (Hanning et al., 2010).

Profilin 3, on the other hand, is an actin-binding protein essential for cytoskeletal dynamics in plants. Its conserved structure across different plant taxa renders it a potent cross-reactive allergen. Research indicates that profilins, including Profilin 3, are responsible for triggering up to 20% of pollen allergies, with Profilin 3 being one of the most significant contributors to this phenomenon. The structural similarities of Profilin 3 across various species enhance its potential to elicit allergic reactions in individuals sensitized to multiple plant allergens (*PFN3 Profilin 3 [Homo Sapiens (Human)] - Gene - NCBI, 2025*).

Together, HSP 70 and Profilin 3 represent critical targets for allergy management strategies aimed at reducing the allergenic impact of *Populus tomentosa* pollen. By suppressing the expression levels of these pan-allergens, it may be possible to alleviate potential allergic symptoms. According to the allergen threshold theory, allergic symptoms arise only when allergen exposure exceeds a certain threshold level (AAH, 2019). Thus, reducing allergen levels below this threshold can prevent or alleviate allergic responses. By lowering the expression of major allergens like HSP70 and Profilin 3, the total allergen load released into the environment is decreased, which can keep exposure below the symptomatic threshold for many allergic individuals. This approach not only helps mitigate symptoms but also improves quality of life by addressing the molecular drivers of pollen allergenicity, making it an effective strategy for managing pollen allergies.

We seek an innovative solution that demarcates itself from previous solutions to pollen allergies that focused on treatments for human symptoms or interventions for the trees themselves. Conventional human-centered approaches, such as using antihistamines, require daily medication to manage symptoms and can cause undesirable side effects like drowsiness (Stephanie, 2025). Similarly, immunotherapy methods, such as allergy shots, demands a long-term commitment of regular injections to build tolerance to allergens. Interventions that target the trees themselves, such as gibberellic acid, is a commonly injected medicine for trees; however, it causes elongated and weak stems, making plants more prone to lodging (Agriplex, 2023). To circumvent these drawbacks, our project employs a targeted, transient biotechnological approach at the source: RNA interference (RNAi).

RNA interference (RNAi), a natural cellular process that uses small interfering RNA (siRNA) to silence the expression of specific genes—in this case, HSP70 and Profilin 3 (Nih.gov, 2017). The siRNA is delivered using a sophisticated nanocarrier system of polyethylenimine-coated dendrimers (PEI-CDs). Dendrimers are highly branched, nanosized polymers that can be engineered to carry a significant payload of siRNA, while the polyethylenimine (PEI) coating provides a positive charge, facilitating efficient uptake into the plant’s cells (Cao, Zhao and Zeng, 2019). Crucially, this method offers a targeted and transient solution. Unlike permanent genetic modification, RNAi only temporarily suppresses allergen production for a single pollen season and is biodegradable, leaving no permanent trace in the tree or environment. It also poses no risk of creating weaker tree structures, as it merely interrupts the production of specific proteins rather than altering growth hormones, making it a precise and harmless alternative to previous methods.

Our project aims to utilize leaf and trunk injection methods to deliver siRNA-loaded polyethylenimine-coated dendrimers (PEI-CDs) into *Populus tomentosa* trees. This biotechnological intervention seeks to suppress the expression levels of key allergens such as Heat Shock Protein 70 (HSP 70) and Profilin 3 (PFN 3), thereby potentially reducing the allergenic impact of these trees on susceptible populations. Verification was completed by performing agarose gel electrophoresis, showing that RNA was fully loaded on the positively charged nanomaterials, and RT-qPCR showed a successful suppression of pan-allergens. By addressing the root cause of pollen allergies through advanced genetic techniques, our research could pave the way for more sustainable and effective allergy prevention strategies in urban settings like Beijing and beyond.

## 2 Materials and Methods

Specific regions of the target genes (HSP70 and Profilin 3) were identified using the NCBI database. For genes with multiple transcripts, the consensus coding sequence (CCDS) was selected to target all functional isoforms. Initial siRNA sequences were designed using an online RNAi Designer tool (Thermo Fisher Scientific)^1^.

To ensure high efficacy and specificity, the selection process adhered to established siRNA design principles. Candidates were required to have a GC content between 30-55%, avoid known SNP sites and repetitive elements, and exhibit asymmetric thermodynamic stability to promote correct strand loading into the RNA-induced silencing complex (RISC). Furthermore, a stringent BLASTN analysis against the *Populus trichocarpa* transcriptome was performed to select sequences with minimal homology (>3 mismatches) to non-target genes, thereby mitigating potential off-target effects.

To expand the specificity analysis, the cross-species efficacy of the selected siRNAs for *Nicotiana tabacum*, *Osmanthus fragrans*, and *Gardenia jasminoides*, the percentage identity and continuous sequence alignment between the siRNA target sites and the homologous HSP70 and Profilin 3 genes in all three species were rigorously evaluated. The siRNA sequences, designed against the *Populus trichocarpa* transcriptome reference sequences were aligned to the coding sequences of the target genes from each species using the NCBI Nucleotide BLAST (BLASTN) suite with standard parameters (Altschul et al., 1990). A minimum sequence identity threshold of 21/21 (100%) or 20/21 (≥95%) nucleotides within the 21-nt siRNA guide strand core targeting region was required for inclusion, as even a single central mismatch can severely abrogate RNAi silencing efficiency (Birmingham et al., 2006; Elbashir et al., 2001). This stringent in silico analysis confirmed that the selected siRNAs possessed the high degree of homology necessary for effective gene silencing across the studied plant species prior to experimental validation.

Given the inherent unpredictability of siRNA efficiency due to factors like local mRNA secondary structure, three distinct siRNA sequences per target gene, each with a high prediction score (≥4.5 stars) and targeting regions separated by more than 25 base pairs, were selected for chemical synthesis. These candidates were then experimentally screened for silencing efficiency via RT-qPCR in a protoplast model. The single most effective siRNA for each target gene was chosen for nanomaterial loading and further functional studies.

Two oligonucleotides, representing the complimentary sense and antisense strands of each desired siRNA respectively, were synthesized:

Oligo sequence for siRNA synthesis:

siHSP70-1-SS: CCUUCAAGGUCAUCGAGAAGG,

siHSP70-1-AS: UUCUCGAUGACCUUGAAGGGG;

siHSP70-2-SS CCUACGGUCUUGACAAGAAGC,

siHSP70-2-AS: UUCUUGUCAAGACCGUAGGCA;

siHSP70-3-SS: CCAUGUACCUCACCAAGAUGC,

siHSP70-3-AS: AUCUUGGUGAGGAUCAUGGAG;

siPFN3-1-SS: GGUUGCUGCUAUCAUGAAATT,

siPFN3-1-AS: UUUCAUGAUAGCAGCAACCTT;

siPFN2-2-SS: CAAAGUACAUGGUGAUCCAGG,

siPFN2-2-AS: UGGAUCACCAUGUACUUUGUG;

siPFN3-3-SS: GGAGCUGUGAUUCGUGGAAAG,

siPFN3-3-AS: UUCCACGAAUCACAGCUCCGG. *

### 2.1. Microbial Cultivation

#### 2.1.1 Bacterial Culture

We labeled 12 culture tubes (2 each for pA7-GFP, PA7-HSP70, PA7-PFN3, pPICZαA, pPICZαA-HSP70, pPICZαA-PFN3) and added 2 mL LB media to each. We added 2 μL ampicillin to pA7 tubes and 1 μL blasticidin to pPICZαA tubes, followed by 100 μL bacterial culture. We sealed the tubes and incubated them at 37°C, 220 rpm for 14 hours.

For larger cultures, we prepared 6 tubes (2 per construct) with 6 mL LB broth, 2 μL ampicillin, and ∼130 μL bacterial colonies, incubating at 37°C, 220 rpm for 19 hours.

### 2.2. Molecular Cloning

#### 2.2.1 Plasmid Extraction

We labeled 12 microcentrifuge tubes and pipetted 1.2-1.5 mL bacterial culture into each. After centrifuging at 12,000 rpm for 1 minute, we removed the supernatant and added 250 μL Solution I to each tube. We added 250 μL Solution II, inverted gently 8 times, then added 350 μL Solution III within 5 minutes and inverted again. After centrifuging at 12,000 rpm for 10 minutes, we transferred the supernatant to new tubes, added ethanol, and loaded onto spin columns. We washed twice with 750 μL Wash Solution before eluting with 50 μL Elution Buffer.

#### 2.2.2 PCR Amplification

We set up reactions in 8 tubes containing 25 μL PCR mix, 2 μL each of forward/reverse primers, 2 μL DNA template, and 19 μL water. We ran PCR with: 95°C for 5 minutes; 40 cycles of 95°C/30s, 60°C/30s, 72°C/2min; final extension at 72°C for 10 minutes.

#### 2.2.3 Enzyme Digestion

We digested plasmids with EcoRI/SalI (pPICZαA) or XhoI/SalI (PA7) in buffer at 37°C for 30 minutes.

#### 2.2.4 Gel Electrophoresis

We prepared 1.2% agarose gels in TAE buffer with GelRed, loaded samples with loading buffer, and ran at 180V for 15 minutes. After UV visualization, we excised bands, dissolved them in Buffer B2, and purified using spin columns.

### 2.3. Preparation and Transformation of Plant Protoplasts

#### 2.3.1. Plant Material and Protoplast Isolation

Protoplasts were isolated from the rosette leaves of 4-week-old Arabidopsis thaliana - ecotype Col-0 - plants grown under a 16/8 h light/dark cycle at 22°C. Fully expanded leaves were harvested, sliced into 0.5-1 mm strips with a sharp razor blade, and immediately transferred into a plasmolysis solution (0.6 M mannitol, 20 mM MES, pH 5.7) for 30 minutes.

The enzyme solution was prepared by dissolving 1.5% (w/v) cellulase R-10 (Yakult Pharmaceutical) and 0.4% (w/v) macerozyme R-10 (Yakult Pharmaceutical) in a digestion buffer containing 0.4 M mannitol, 20 mM KCl, 20 mM MES (pH 5.7), and 10 mM CaCl . The solution was incubated at 55°C for 10 minutes, cooled to room temperature, and then supplemented with 0.1% (w/v) bovine serum albumin (BSA) and 5 mM β-mercaptoethanol. The solution was filter-sterilized through a 0.22 μm membrane.

The plasmolysis solution was removed from the leaf strips, which were then submerged in the enzyme solution (10 mL per 1 g of leaf tissue). Digestion was carried out in the dark for 3 hours with gentle shaking at 40 rpm.

#### 2.3.2. Protoplast Purification and Transformation

The digestate was gently passed through a 70 μm nylon mesh filter to remove undigested debris. The filtrate was transferred to a 15 mL conical tube and an equal volume of W5 solution (154 mM NaCl, 125 mM CaCl , 5 mM KCl, 5 mM glucose, 2 mM MES, pH 5.7) was added to dilute the enzymes. Protoplasts were pelleted by centrifugation at 100 x g for 3 minutes at 4°C. The supernatant was aspirated, and the pellet resuspended in W5 solution. The protoplasts were counted using a hemocytometer and diluted to a density of 2 x 10 cells/mL in W5 solution. The cells were iced for 30 minutes.

For transformation, approximately 2 x 10 protoplasts (in 100 μL) were aliquoted into a 2 mL microcentrifuge tube. 10 μg of plasmid DNA (or an equivalent amount of PEI-CDs@siRNA nanocomposite) and 110 μL of freshly prepared PEG solution (40% (w/v) PEG 4000, 0.2 M mannitol, 0.1 M CaCl ) were added. The mixture was mixed by inversion and incubated at room temperature for 15 minutes. The transformation was stopped by adding 440 μL of W5 solution while mixing. The transformed protoplasts were pelleted at 100 x g for 3 minutes, the supernatant was removed, and the pellet resuspended in 1 mL of WI solution (0.5 M mannitol, 20 mM KCl, 4 mM MES, pH 5.7). The cells were transferred to a 12-well plate and incubated in the dark at 22°C for 16-48 hours before subsequent analysis.

### 2.4. Preparation of Nanomaterials

#### 2.4.1 Carbon Dot Synthesis

We combined glycerol and polyethyleneimine, microwaving for 2 minutes (3 cycles). After cooling, we added water and sonicated for 30 minutes.

#### 2.4.2 Purification

We dialyzed the product against deionized water using dialysis bags, changing buffer periodically. We then freeze-dried the purified carbon dots for storage.

### 2.5. Synthesis of PEI-CDs@siRNA Nanocomposites

#### 2.5.1 Evaluation of Loading Efficiency

We centrifuged mixtures PEI-CDs@siRNA at mass ratios of 1:0, 1:1, 1:2, 1:5, 1:10, 1:20, 1:50 and 1:100 respectively and applied electrostatic adsorption incubation for 30 minutes to ensure full integration. Then, we conducted the electrophoretic mobility shift essay for all mass ratios sets on 150V for 30 minutes to validify effective loading ratios.

#### 2.5.2 Loading of siRNA onto PEI-CDs

We centrifuged mixtures PEI-CDs@siRNA at mass ratios of 1:100 and applied electrostatic adsorption incubation for 30 minutes to ensure full integration, procuring the product nanocomposite.

### 2.6. Method of Administration

*Nicotiana tabacum* were grown under a 16/8 hour light/dark cycle at 24°C and 60% relative humidity for 6 weeks prior to injection.

*Osmanthus fragrans* and *Gardenia jasminoides* samplings were grown a under a 16/8 hour light/dark cycle at a daytime temperature of 26 ± 2 °C, nighttime temperature of 22 ± 2 °C, and relative humidity at 65-75% for 16 weeks prior to injection.

#### 2.6.1 Leaf Injection

We injected PEI-CDs@siRNA nanocomposites of 100:1 ratio into the lower epidermis of Nicotiana tabacum leaves using sterile syringes.

#### 2.6.2 Trunk Injection

We drilled 10 mm holes in *Osmanthus fragrans* and *Gardenia jasminoides* trunks and injected nanocomposite solution at 0.1 mL/5 min intervals before sealing.

### 2.7. Confirmatory Gene Silencing

#### 2.7.1 RNA Extraction

After 16 hours, we extracted the leaves, homogenized leaf tissue in liquid nitrogen, lysed in buffer, and purified RNA using the Spectrum™ Plant Total RNA Kit (Millipore Sigma) with on-column DNase I digestion.

#### 2.7.2 RT-qPCR

We synthesized cDNA from 1 µg RNA using the High-Capacity cDNA Reverse Transcription Kit (Thermo Fisher Scientific) and performed qPCR with gene-specific primers using PowerUp™ SYBR™ Green Master Mix (Thermo Fisher Scientific) on a QuantStudio™ 3 system. We calculated relative expression using the 2^(-ΔΔCt) method with β-actin normalization.

## 3. Results

siRNA sequences targeting two candidate allergenic proteins (Potri.010G205700.1, HSP70 family; Potri.003G047700, Profilin family) were loaded onto PEI-modified carbon dots (PEI-CDs) by electrostatic adsorption to form PEI-CDs@siRNA nanocomposites. Experiments were performed across three biological scales (Arabidopsis mesophyll protoplasts, Nicotiana tabacum leaf injections, and trunk injections into woody species) to evaluate delivery efficiency and target mRNA modulation in different tissue and organism contexts.

This approach allows a thorough evaluation of the efficiency of this RNAi structure in reducing allergenic protein expression in biological systems of different sizes and types.

### 3.1 Optimal Formulation of PEI-CDs@siRNA

Negatively charged siRNA was loaded onto synthesized PEI-CDs compounds through electrostatic adsorption, resulting in the formation of the PEI-CDs@siRNA nanocomposite. To evaluate the loading efficiency of siRNA, siRNA and PEI-CDs was combined in different mass ratios: 1:0, 1:1, 1:2, 1:5, 1:10, 1:20, 1:50 and 1:100 respectively. The mixture was centrifugated and incubated for 30 minutes for full integration.

Agarose gel electrophoresis across a PEI-CDs:siRNA mass ratio series (1:0 → 1:100) showed free siRNA bands at lower ratios and loss of free siRNA migration at 1:50 and 1:100, consistent with near-complete complexation at these ratios (Figure 1). Based on these data we selected a 1:100 mass ratio for downstream experiments.

**Figure 1.**
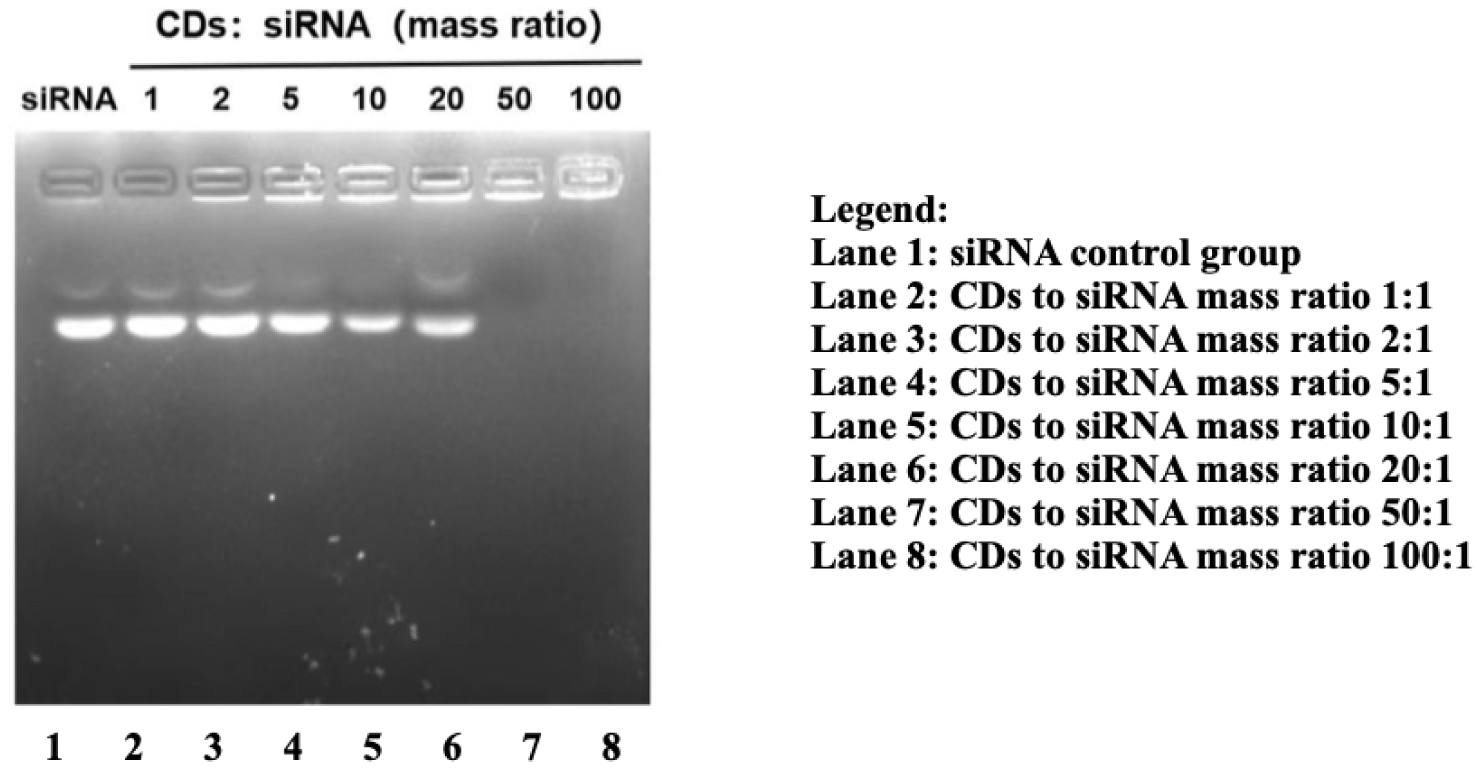
Electrophoretic Mobility Shift Assay for Analysis of CDs’ Loading of siRNA.

### 3.2 Delivery of siRNA into Arabidopsis Protoplasts via Transformation

*Arabidopsis thaliana* protoplasts were transformed with 12 siRNA variants (three sequences per target, tested with and without PEI-CD encapsulation: siHSP-1, 2 & 3 with and without PEI-CDs encapsulation, siProfilin-1, 2 & 3 with and without PEI-CDs encapsulation) as exogenous siRNA input to evaluate sequence- and carrier-dependent effects on target transcript levels. Transformation was carried out with the Thermofisher transformation kit and solutions, following given instructions.

Using the two-step technique, Real-time quantitative reverse transcription polymerase chain reaction (RT-qPCR) was conducted on the RNA extracted from the plant tissues. The ΔΔCt relative quantification method was used to analyze results attained with β-actin as the reference gene. The two-step method for RT-qPCR was selected, involving a reverse transcription and a real-time PCR, indicating the real-time monitoring of the amplification of the genes of interest through examining corresponding fluorescent markers. Assays included 3 biological replicates (reported in figure legends).

#### 3.2.1 Data Analysis

Results from this experiment aim to demonstrate the difference in silencing effectiveness between the three different siRNAs developed for each protein. In this examination, the control group did not receive any siRNA, therefore establishing a baseline for comparison against treatment groups. Figure 2 refers to the 3 different siRNA sequences designed to assess their compatibility with the targeted gene. For PFN3, the most pronounced changes were observed for one Profilin siRNA (PFN 3-A), which produced a mean reduction to ∼0.50-fold of control (mean ± SD;). The other two PFN-targeting siRNA (B and C) were not successful, since their mRNA levels remained mostly unchanged.

**Figure 2.**
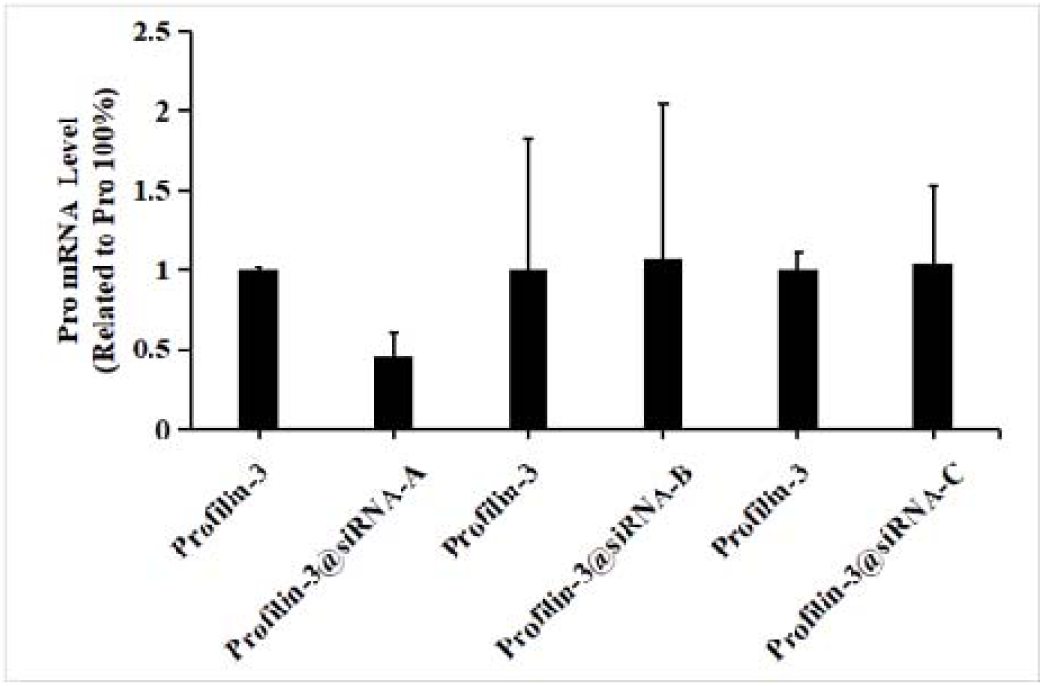
Profilin 3 mRNA transcription level in treated and untreated *Arabidopsis thaliana* protoplasts

As shown in Figure 3, the three siHSP70 sets showcased great discrepancies. For HSP70, siHSP70-2 produced the largest mean decrease (∼80% reduction in relative transcript; mean ± SD;), while siHSP70-3 produced a more modest mean decrease (∼30% reduction). siHSP70-1 showed minimum influence on the mRNA transcription levels. Contradictory to expectation, the value of HSP 70 mRNA raised slightly after the application of the therapy. Yet, the scale of increase is low, suggesting the change constitutes as a normal fluctuation rather than a causal relationship.

**Figure 3.**
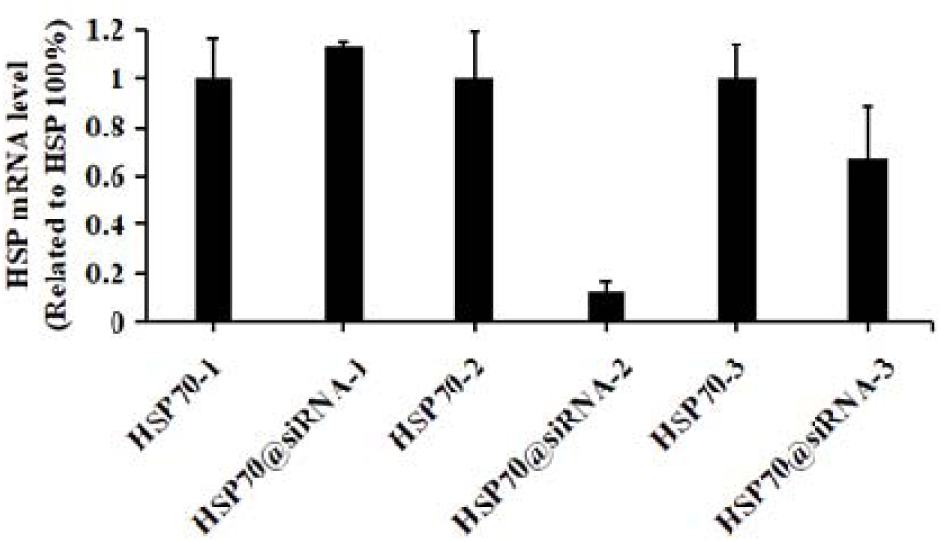
HSP 70 mRNA transcription level in treated and untreated *Arabidopsis thaliana* protoplasts

These protoplast data indicate sequence-dependent knockdown potential but also show notable variability across replicates. Given the limited sample sizes in the current dataset, these observations are preliminary; statistical details and exact p-values are included in figure legends and Methods. Follow-up experiments with additional biological replicates and independent siRNA sequences are planned to confirm these findings.

### 3.3 Delivery of siRNA into *Nicotiana tabacum* via Leaf Injection

siRNA was delivered into Nicotiana tabacum leaves using an insulin syringe without the needle to aspirate and inject the liquid. The plant was chosen to verify the utility of the construct because, as a model organism, it enables greater experimental control and a deeper understanding of the system. Additionally, its representation of herbaceous systems helps broaden the treatment’s scope of applicability. During the injection, the leaves were turned upside-down, and the liquid was pushed through the syringe on to the leaf, ensuring that venules were bypassed. Each treatment was tested in 3 independent trials (biological replicates).

#### 3.3.1 Data Analysis

The results of our RT-qPCR efficiency testing experiments showed that the uptake of siRNAs by leaf injection was practicable and effective in silencing HSP1/2/3 and profilin2/3. Contradictory to our expectation, profilin-1 did not yield successful results. In fact, instead of negatively regulating the expression, it induced an increase of about 5%. However, according to our preliminary research, it was discovered that usually, an alteration lower than 50% has low statistical significance, therefore the modification created by Profilin-1 can be seen as a fluctuation in the normal mRNA levels in an unmodified plant, thus showing Profilin-siRNA-1 was simply unsuccessful, instead of contradicting other results. Also, it was found that, in protoplasts, HSP-3 was the best siRNA targeting HSP 70, restraining over 70% of HSP 3 production (mean ± SD;). Profilin-2 was the best siRNA targeting Profilin 3, reducing Profilin 3 expression by ∼60% (mean ± SD). This contradicted with the performance of the sequences in the *Arabidopsis* assays, suggesting an influence of different metabolisms and structures on the siRNAs’ compatibility to the target mRNA.

**Figure 4.**
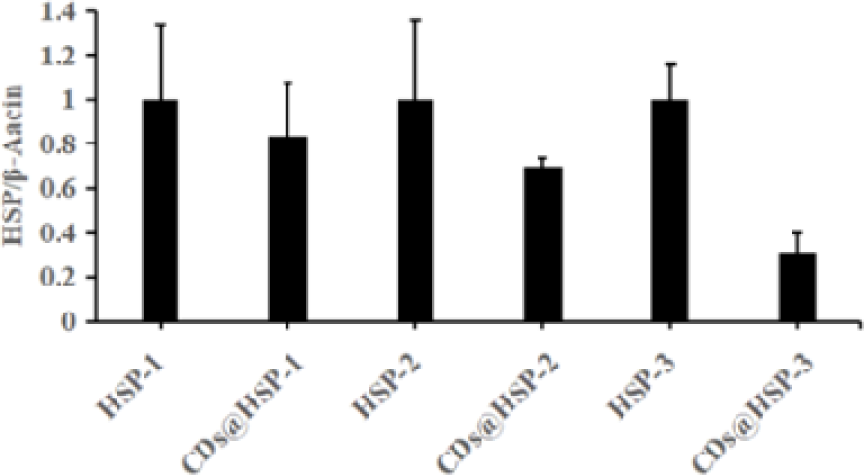
RT-qPCR: HSP 70 in *Nicotiana tabacum* leaves.

**Figure 5.**
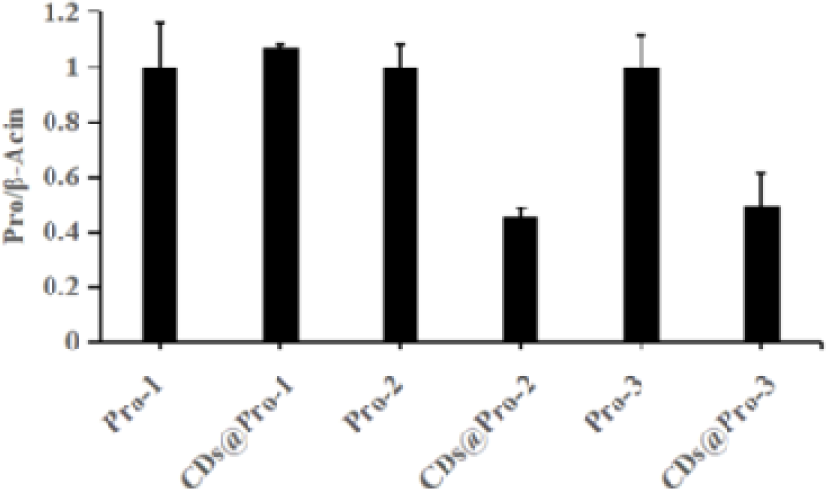
RT-qPCR: Profilin 3 in *Nicotiana tabacum* leaves

### 3.4 Delivery of siRNA into *Gardenia jasminoides and Osmanthus fragrans* via Trunk Injection

Exogenous siRNA molecules were delivered via trunk injection into Osmanthus fragrans, two agronomically significant woody plant, which were chosen as, similar to *Populus tomentosa* trees, it also possess woody trunk tissues. The tested plant averaged 1.5m in height and 10cm in diameter. Employing an electric drill, holes of 6mm diameter and 4 mm depth were drilled at a 45-degree angle beneath main boughs of each tree. The 12 different variations of composites are loaded into 12 separate sterile, 1mL insulin syringe (without needles). Through this device, the solution was gently, slowly injected into the trunk through the holes. Untreated trees were used as controls with 1mL of Diethyl Pyrocarbonate water injected with the same method. The plants were injected once within a single hour, then kept at room temperature under light.

16 hours after the injection, leaf samples at sites distant from the injection point were collected and marked. Collected samples were submersed in liquid nitrogen and ground into powder using separate mortars and pestles. RNA extraction was performed on 30mg of powdered leaf tissue per sample to isolate the targeted HSP 70 and Profilin 3 mRNA to assess the change of target mRNA after siRNA therapy suppression. RT-qCPR was then performed on extracted RNA samples. 3 technical replicates were conducted for each protein.

#### 3.4.1 Data Analysis

The analysis of targeted HSP 70 and Profilin 3 mRNA concentration in the treated and untreated samples using different targeting siRNA sequences successfully displayed a statistical difference in the suppression rate.

PEI-CDs-encapsulated HSP70 siRNAs produced modest mean reductions in HSP70 transcript levels across treatments (approximately 14%, 35%, and 44% mean reductions for sequences HSP70-1/2/3 respectively; mean ± SD for each). All naked (unencapsulated) siRNAs produced no consistent knockdown, consistent with poor uptake in the absence of carrier. A potential explanation of such phenomenon is the presence of a negative charge in the cell wall that hindered the entrance of extraneous RNA via repulsive forces, justifying the necessity for nanocomposite encapsulation when considering siRNA inhibition in plants.

**Figure 6.**
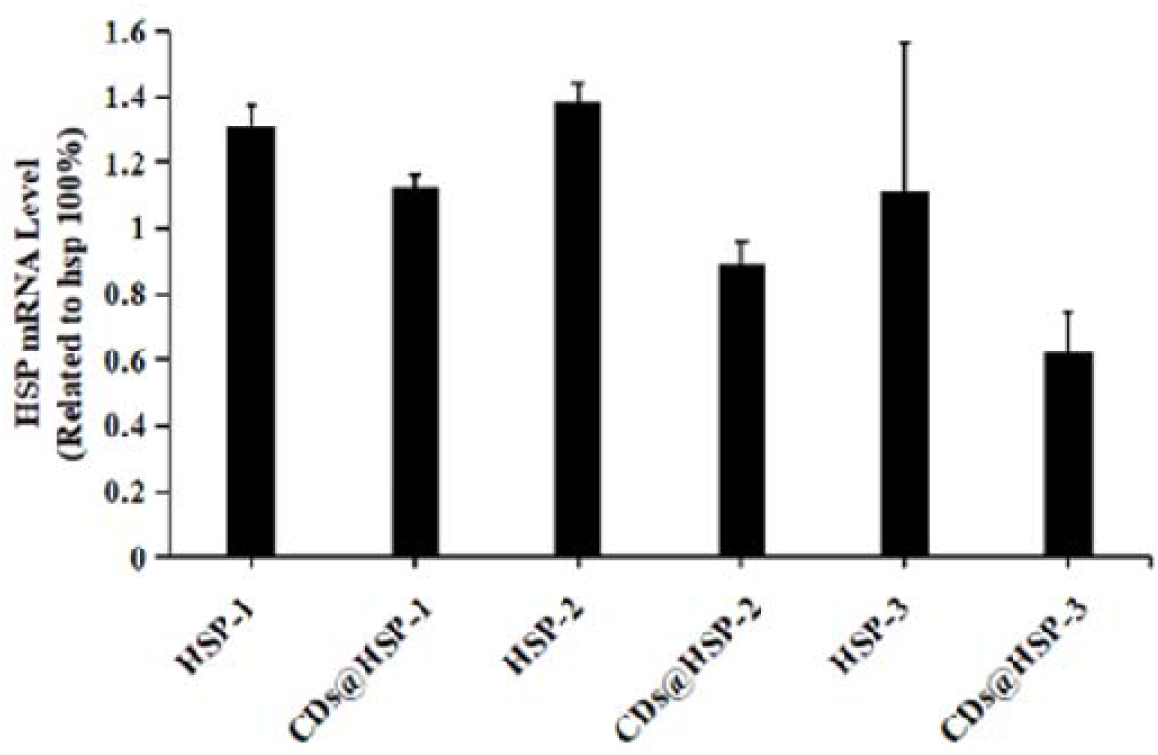
RT-qPCR: Trunk injection levels of HSP 70 in *Osmanthus fragrans* leaves.

Profilin-targeting siRNAs yielded inconsistent outcomes in trunk injections: mean transcript changes were +6%, +29%, and +54% for Profilin-1/2/3. These increases are inconsistent with the knockdown observed in other systems and with expected outcomes. This observation, combined with results from previous methods, suggests an overall inconsistent and sometimes ineffectual repression of targeted mRNAs by the designed siRNA sequence. Possible technical explanations include sequence-dependent off-target effects, delivery inefficiency, or biological feedback regulation. We note that no scrambled (non-targeting) siRNA or vehicle (PEI-CDs-only) controls were included in the present dataset. Therefore, we cannot definitively attribute the observed increases to sequence-specific effects. Follow-up experiments will include scrambled controls, vehicle controls, redesigned siRNAs, and protein-level validation to clarify these outcomes.

**Figure 7.**
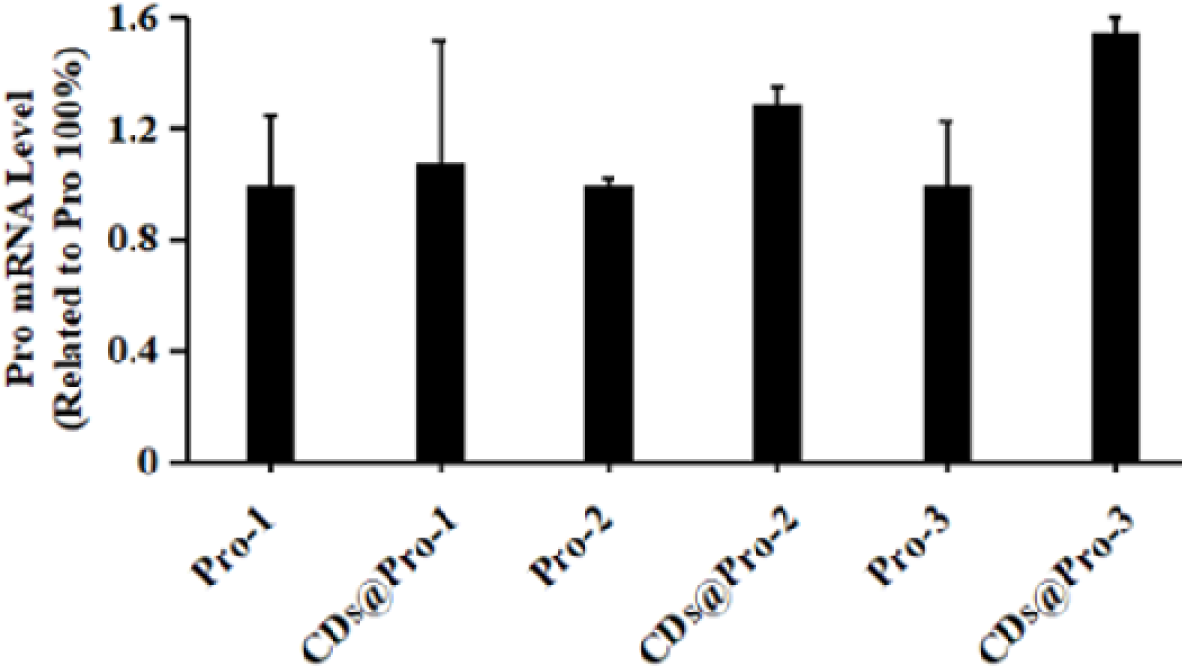
RT-qPCR: Trunk injection levels of Profilin 3 in *Osmanthus fragrans* leaves

## 4. Discussion

The results from rt-qPCR demonstrate the ability of the developed PEI-CDs-siRNA construct to suppress targeted mRNA transcripts of putative allergens Hsp70 and Profilin 3 by a 14.10% to 87.29% reduction rate in several plant systems (i.e., *Arabidopsis thaliana*, *Nicotiana tabacum*, *Osmanthus fragrans*) and delivery approaches (i.e., protoplast transformation, leaf injection, trunk injection) studied. Moreover, loading siRNA into PEI-CDs nanomaterials also proved to be efficient at ratios 1:50 and 1:100. These data hence suggest a possibility in the utilization of the PEI-CDs-siRNA complex for the inhibition of plant allergens.

Previous research has extensively explored the mechanism, engineering, and applications of RNAi-mediated inhibition of allergens in edible agricultural crops, including tomatoes, apples, and peanuts, yielding significant results. Inhibition of minor allergen Lyc e 3 in tomatoes L. esculentum var. Microtom through RNAi was able to reduce allergenic protein to below detection levels, with a 10-100 fold higher protein concentration level to induce the same histamine release level as wild-type lines (Le et al., 2006). Similarly, Mal d 1 in apple cultivar E94 observed a minimum of 10-fold reduction in allergen accumulation according to immunoblotting and prick-to-prick tests (Gilissen et al., 2005). However, similar to our study, only leave samples were tested due to the long regeneration time of the targeted tissue. Thus, the feasibility of RNAi to reduce allergens in the fruit itself requires further confirmation. Ara h 2 and Ara h 6 were documented to be completely silenced in some peanut lines through quantitative immunoblotting with no observed negative effects on seedling growth (Chu et al., 2008). Still, a limited number of RNAi approaches directed towards non-food sources have been reported.

As such, our study complements the current focus on ingested allergens by discussing applications of RNAi on inhaled allergens such as plant pollen, confirming the mRNA suppression potential of the PEI-CDs-siRNA strategy.

Moreover, despite the current prevalence of delivery tools such as agrobacterium-mediated and gene gun transformed transgenic plants, these systems limit RNAi’s full potential in plants with regard to various aspects. Agroinfiltration, for one, though it provides high siRNA expression levels, requires DNA to be encoded in a viral vector and transported into the plants through the pathogenic agrobacterium. This presents two key limitations on its wide-range application:

Firstly, it is only useful in a limited scope of susceptible species, with recognized instances of off-target effects in some studies (Kodama and Komamine, 2011; Xu et al., 2006); in addition, as these vectors are integrated into the hosts’ genomes, it creates transgenic plants that may inflict public concern. Meanwhile, biolistic methods exhibited lower transformation efficacies. For these reasons, many studies, such as that of Dalakouras et al. (2018) and Wise et al. (2022), are experimenting with multiple delivery strategies for introducing extraneous siRNA and dsRNA into plant systems, in seeking alternative RNAi cargoes that can bypass the rigid plant cell walls, be widely applied to plants while also avoiding damage. Situated in such a background, our study provides some insights into how the same set of siRNA sequences may function differently through different delivery approaches.

Nonetheless, though our identification of the relative inhibition rate of the sequences on the mRNA level is a critical first step to evaluating the success of the approach, additional studies and assays are required to conclusively prove the effectiveness of the treatment.

For example, the relationship between reduced mRNA transcription and the successful suppression of protein expression remains complex and requires further exploration. Thus, one further step can be the evaluation of protein expression levels, especially in the targeted pollen tissues. Through sampling pollen in wild-type plants and RNAi plants and subjecting attained proteins to western blotting, a comparative analysis of protein expression levels can assist researchers in understanding how siRNA suppression effects can be sustained or counteracted in translation processes. As compensatory feedback loops, exemplified by the regulation mechanism of Hsp 70 in Saccharomyces cerevisiae (Garde et al., 2024), may upregulate transcription in response to a perceived deficiency to maintain homeostasis. The fact that some studies (Peyman Aghaie & Hosseini, 2020; Anaraki et al., 2018) have reported a central role of 70kDa heat shock proteins in plants’ abilities to adapt to drought and other types of environmental stress specifically underscores the high possibility that plant systems may upregulate these allergenic proteins.

It is also critical to note that the current idea on the reduction of immunological responses in humans is only theoretical. To further confirm the effectiveness of these siRNA sequences. To this end, future studies could consider collecting patient sera to conduct immunoblotting and histamine release assays. For example, binding rates of Hsp 70 and Profilin 3 to specific IgE antibodies in wild-type and RNAi plants and cross reactivity, as well as histamine release level in basophil and mast cells in patients after being exposed to the allergens, could be measured. Such assays would not only provide valuable data for the verification of the isoforms’ allergenicity but could also help make a conclusive statement about the ability of this approach to reduce allergenic reactions in the human body.

Lastly, the influences of the siRNA injection on the plant health and natural growth remain unknown. The pollen tissue, for instance, as one major tissue influenced by the allergen suppression, also plays a significant role in the reproduction of plants; the fertility of the RNAi plants needs to be confirmed through germination assays before it can be applied in real-life environments. Therefore, the role of the specific allergenic isoform targeted by the developed siRNA sequences still needs to undergo thorough functionality analysis to determine if its role in the plant system can be substituted. Specifically, the impact that siRNA exerts on photosynthetic efficiency, plant growth rate, reproductive fitness, and overall physiological health should be evaluated in future research, as such assessments are needed to ensure that siRNA therapeutics minimize allergenicity but do not impose unintended harm upon the treated plants.

Overall, the discussed treatment does provide insights into the potential usefulness of siRNA in allergy prevention treatments, indicating that research into PEI-CDs mediated RNAi is a promising and innovative strategy for obtaining hypoallergenic plant pollens. Yet, the dataset still has clear limitations: (1) several comparisons were performed with small biological sample sizes; (2) though a baseline control group was established through injecting Diethyl Pyrocarbonate water, negative controls, such as non-targeting siRNA sequences or PEI-CDs-only solutions, were not included. Therefore, its functionality, applicability, and scalability still remain undetermined. To strengthen the conclusions, follow-up experiments and future studies could include: (1) addition of scrambled and vehicle controls run in parallel with all siRNAs; (2) redesign and testing of additional independent siRNAs per gene; (3) siRNA uptake assays (fluorescently labelled siRNA); and (4) protein-level validation and immunological assays.

## Author Contributions

All authors contributed equally to this work. ZL, BR, KY, PZ, and SZ jointly conceptualized the study, designed the methodology, conducted the experiments, analyzed the data, and wrote the manuscript. All authors reviewed and approved the final version of the manuscript.

## Acknowledgments

The authors would like to express their sincere gratitude to Dr. Zhu Ke for his invaluable guidance throughout the study. We appreciate the China Academy of Chinese Medical Sciences and China Agricultural University for providing the laboratory.

## Footnotes

1 https://rnaidesigner.thermofisher.com

